# Multiple Kernel Learning approach for Medical Image Analysis

**DOI:** 10.1101/121509

**Authors:** Nisar Wani, Khalid Raza

## Abstract

Computer aided diagnosis is gradually making its way into the domain of medical research and clinical diagnosis. With field of radiology and diagnostic imaging producing petabytes of image data. Machine learning tools, particularly kernel based algorithms seem to be an obvious choice to process and analyze this high dimensional and heterogeneous data. In this chapter, after presenting a breif description about nature of medical images, image features and basics in machine learning and kernel methods, we present the application of multiple kernel learning algorithms for medical image analysis.

## 1 Introduction

Biological systems are made up of many subsystems. These subsystems, also called organs are very complex and house sophisticated machinery to carry out various physiological functions. The organization of human body can be viewed as an integrated system where each specific organ contributes in a special manner, both in terms of anatomical characteristics and physiological functions. A brief description of some important sub systems is given in Table 1.

**Table 1:** Some important subsystems of human body.

| Name | Function |
| --- | --- |
| Visual System | Performs the physiological function of vision. Image information is focused onto the retina, transduced as neuronal signals and transmitted to visual cortex. The visual cortex finally decodes the received signal |
| Digestive System | Is essential for the intake, breakdown and absorption of essential nutrients, which the body uses for energy, growth and cell repair. |
| Cardio Vascular System | Performs the primary task of transporting the essential nutrients to almost all cells of the body via its arterial network and is also vital to replenishing the oxygen content of the blood through the pulmonary system via lungs. |
| Nervous System | Coordinates the activity of muscles; controls and monitors organ functions and responses, transmits and processes data received from other sensory organs as neuronal signals.<br>Brain and spinal-cord are very important subsystems of the nervous system called “Central Nervous System” that act as control centres for handling almost all sorts of neuronal communication within the human body. |
| Musculoskeletal System | The system gives shape, support, stability, and overall organization to the body. The system is made up bones, muscles, cartilages, and other connective tissues. It binds tissues and organs together and is therefore responsible for giving them special anatomical characteristics viz. shape, density and texture etc. |

All these subsystems exhibit certain anatomical characteristics and play vital role in many physiological processes. The anatomical characteristics are manifested as shape or texture patterns that demonstrate the integrity of the system itself and well being of individual organs. However, the physiological functions are very complex phenomena that include hormonal stimulation and control, information gathering and processing, internal and external communication, responses in the form of mechanical actions, biochemical reactions and electrical signals which reflect their nature and activity. Any deviation from the normal physiology affects health, performance and overall functionality of the system thus giving rise to a wide range of pathological conditions associated with physiological functions and anatomical features, which are quite contrary to the corresponding normal patterns.

With a good understanding of the system of interest, capturing its anatomical and physiological information is essential for making efficient clinical decisions encompassing a wide range of medical complications that may arise during the lifetime of a biological being. Medical imaging is one such non–invasive procedure that allows medical practitioners to study the internal state of the organs from both anatomical and physiological perspectives to obtain important clues about a medical condition without actually opening those organs or without using any other intrusive method. Medical imaging (Digital X-Ray, MRI, CT scan, PET scan etc.) has emerged as an important discipline and a tool to guide the diagnosis of deformed and dysfunctional organs, such as detecting bone fractures, calcification within organs, organ enlargement and all the way to advanced brain tumours, cardiac malfunctions and malignant growths in other vital organs.

This tutorial is organised into 8 sections and many subsections. After presenting the introduction in section 1, we present nature of biomedical images in section 2. Section 3 is devoted to image descriptors, section 4 covers computer aided diagnosis. In section 5 and 6 we present basic concepts in machine learning, kernel methods and multiple kernel learning model. Section 7 covers the application of MKL to mammographic data and in section 8 we conclude the chapter.

## 2 Nature and Characterisitcs of Biomedical Images

This section covers the basic imaging modalities used in diagnostic imaging, their characterisitcs and applications.

The aim of biomedical image analysis is to extract valuable information from medical images so as to understand the state of a system under investigation. A number of imaging procedures, invasive and non-invasive, active and passive, have been developed to capture information that is essential for understanding a pathological condition (Rangayyan, 2004). Depending upon the nature of the pathological condition and the composition, structure, function and location of different organs, following imaging techniques are used by modern diagnostic imaging: -

1. **Thermography**: Infrared imaging is a passive, non-invasive imaging method used to detect and locate temperature distribution as a result of some pathological condition while analyzing physiological functions related to thermal homeostasis of the body. Infrared thermography’s use is based on experiences, e.g. certain tumour types, such as breast cancer are expected to be highly vascularised and therefore could be at a slightly higher temperature compared to their neighbouring tissues (Szentkuti et al., 2011). These temperature differences can be measured using thermal sensors for advanced stage of breast cancer, where thermography has been proven as a potential tool. Some pathologic conditions, especially inflammation lead to hyperthermia (areas with higher amount of infrared emission), while degeneration, reduced muscle activity, poor perfusion, and certain types of tumours may cause hypothermia.
2. **Light Microscopy**: Light microscopy provides a useful magnification of up to x1000 and plays an important role in studying cellular structures under different conditions. However, using live cell fluorescence microscopy, dynamic cellular events can be captured in the image format and subsequently subjected to advanced image analysis.
3. **Electron Microscopy:** Electron microscopes provide a resolving power of the order of ×10^6^ and can be useful in revealing the ultra structures of biological cells and tissues. At 60 *keV* an electron beam with a wave length of 0.005 *nm* attains a resolving power limit of up to 0.003 *nm*. Electron microscopes are available in two variants:

> *TEM* (Transmission Electron Microscope): Transmission electron microscopes are the original form of electron microscopes that apply electron beams to illuminate the target specimen in order to create their images. Spherical aberrations dramatically limit the resolving power of TEMs, but the availability of new generation of aberration correctors can be used to increase their resolution (Williams & Carter, 1996).TEMs find application in cancer research, virology, materials science as well as pollution,nanotechnology and semiconductor research.

> *SEM* (Scanning Electron Microscope): The working principle of SEM resembles TEM in many ways. However the electron beam being used by SEM is more finely focused to scan the specimen surface in a raster fashion. SEM operations are carried out in different modes for detecting diverse signals emitted from the specimen. Pictures having a depth of field up to several *mm* can be obtained, which are helpful in analyzing fibre distribution and realignment during the healing process after injuries.(Unakar et al., 1981)
4. **X-ray imaging**: X-Rays are a form of electromagnetic radiations used as an imaging modality for studying internal structures of the human body. An x-ray sensitive film or a digital detector is placed behind the patient to capture the x-rays as they are passed through the patient’s body. Different tissues within the body show varying absorption levels of x-rays, e.g., dense bones absorb more radiation as compared to soft tissues that allow more to pass through. The resulting variance, in absorption levels of the x-rays, produces a contrast within the image to give a two dimensional representation of target organs of the patient. X-ray radiographic applications include examination of chest to assess lung pathology, x-ray imaging of skeletal system to examine bones, joints, diagnose fractures and dislocation etc. Radiography of abdominal cavity is performed to assess obstruction, free air or free fluid. Also the dental x-rays are used to examine cavities and abscesses(WHO, n.d.).
5. **Mammography**: Mammography is an imaging technique used for diagnostic screening and surveillance imaging of breast cancer. Using low energy x-rays and standardized views of the breast, mammograms can be used to inspect breast tissues for lumps, lesions and calcification. Early mammographic examinations can reveal cancer symptoms to the radiologists which allow early treatment and hence increased survival rates(WHO, n.d.). Some of the application areas of the mammography include: -

- Screening mammography for early detection in case of asymptomatic women.
- Diagnostic mammography for diagnosis of a suspected lesion.
- Checking recurrence of malignancy in women treated for breast cancer using surveillance mammography.
- Applicable in tumour marking and needle localization for obtaining tissue samples from suspicious breast masses.
6. **Computed Tomography:** Computed tomography (CT) utilizes x-ray photons for image production with digital reconstruction. The data is converted into digital format by various algorithms once it is captured by the scanner. For each image acquisition task a slightly different angle is set; scanning the whole body in the process. A two-dimensional pixel, each of which carries a designation of density or attenuation, which makes up an image element, and is represented by Hounsfield unit (HU). Computed Tomography has been extensively performed on Brain (with or without contrast and perfusion study). However, it is not only limited to brain but has wider applications, including CT scan of chest (chest CT), abdominal cavity (CT abdomen), CT urography (kidneys, urinary tract, bladder and ureters), blood vessels (CT angiography), colon (CT colonography), heart (Cardiac CT), and spinal cord (CT myelography).
7. **MRI (Magnetic Resonance Imaging)**: This imaging modality in radiology uses magnetic radiations to visualize the internal organs of the body. It offers near perfect 3D views of internal organs in real-time and extremely good contrast of soft tissues, therefore making the visualization of muscles, joints, brain, spinal cord and other anatomical structures much better (Edelman & Warach, 1993).Compilation of sequences that are an ordered combination of radio frequencies and gradient pulses form the basis of MRI visualizations. This type of design is used to acquire the data image formation. MRI imaging is widely used in Brain imaging, spinal cord, MRI of abdomen (to assess liver, spleen, kidneys etc), MRI of blood vessels, heart, joints, muscles and bone disorders
8. **Nuclear Medicine:** This mode of diagnostic imaging uses a radioactive tracer to obtain images of internal organs. A radioactive isotope is added to an organ specific pharmaceutical to obtain images of the target organ. The tracer is basically a gamma radiation emitter. A crystal detector sensitive to gamma rays is fitted into the camera which can detect the distribution of the tracer as it traverses through various body parts. The captured information is digitized to produce 2D or 3D images on a screen. Modern day gamma cameras are shipped along with a CT as hybrid machines opening new vistas for integrated study of nuclear medicine imaging with CT images (Mettler Jr & Guiberteau, 2011). Unlike x-ray studies that detect anatomical alterations, the altered physiology of different organs of the body can be examined with the help of nuclear medicine imaging.
9. **Single Photon Emission Computed Tomography (SPECT):** SPECT is a nuclear medicine tomographic imaging modality that uses gamma rays to provide 3D imaging of target organs, unlike conventional nuclear medicine that produces plane 2D images. A reconstruction algorithm is then employed to obtain 2D planar sections from the individual scan lines of image projections. SPECT can be used to complement any gamma imaging study, where a true 3D representation is needed, e.g., tumour imaging, infection (leukocyte) imaging, thyroid imaging or bone scintigraphy.
10. **Positron Emission Tomography (PET):** PET is an imaging modality that is used to monitor the physiological processes by visualizing blood flow, metabolism, neurotransmission, and drugs labeled with radio tracers (Ollinger & Fessler, 1997). PET offers time course monitoring of disease processes as they evolve or in response to a specific stimulus. PET also uses the radioactive trace for physiological study of the organs as in nuclear medicine. PET scans are commonly performed to identify malignant tumors by measuring the rate of glucose consumption in different parts of the body, considering the fact that malignant tumors metabolise glucose at a faster rate than benign masses. Other applications of PET include understanding strokes and dementia, tracking chemical neurotransmitters (such as dopamine in Parkinson’s disease).
11. **Ultrasound**: This imaging modality employs sound waves of high-frequency to visualize the internal organs of the body. Ultra sound equipment consists of three important components - a monitor, processor and a transducer. The transducer emits sound waves (echoes) at a certain frequency and the probe is gradually moved over organs of interest, the returning echoes are captured and later on digitized to appear as dots on the monitor. Ultrasonogrpahic images can be obtained in any plane and in real time. Ultrasonogrpahy may be conventional real-time or doppler. Doppler uses doppler shift phenomena for performing vascular studies to monitor blood flow and detect blood clots inside veins. Ultrasound is performed to visualize the anatomy and pathology of the liver and gallbladder, spleen, kidneys, lymph nodes, urinary bladder, reproductive organs, vascular structures etc. Cardio vascular ultrasound including ECG is performed to examine the heart and peripheral blood vessels. Ultrasound is also performed in pregnant women to assess the foetus and other structures within the uterus. A brief description of all imaging modalities has been covered in this section, for further details see Rangayyan, 2004.

## 3 Feature Extraction and Description

This section covers basic concepts in feature extraction and image description. The section covers a brief introduction of various image descriptors used in image processing and computer vision domains. Feature extraction is a process in image processing where information contained in the region of interest (ROI) of an image is represented using a feature vector(Nixon & Aguado, 2012). It is also a dimensionality reduction approach, useful when images are of large sizes and a reduced representation is needed to quickly perform image matching and retrieval.

Common feature extraction techniques include HOG (Histogram of Oriented Gradients), SURF (speeded up robust features), LBP (Local Binary Patterns), wavelets, and colour histograms. These are often combined to solve common problems of object detection and recognition in computer vision, content-based image retrieval such as PACS (picture archiving and communication system), face recognition, and texture classification (OPENCV, n.d.).

### Image Descriptors

For robust analysis of biomedical images, the challenge is to extract features from regions of interest (ROI) that accurately describe image elements, such as intensity, texture and shape i.e. properties that can differentiate different features within images. These features are captured by algorithms commonly called as image descriptors (Nogueira et al., 2016), whose output, a feature vector, describes the content of contagious part of an image. Simple descriptors capture all pixel intensities from an ROI, however, due to high dimensionality of the intensity vector these descriptors are computationally inefficient and lack robustness to image distortions. Few descriptors such as SIFT, LBP, HOG etc. make use of distribution based strategy. The descriptors based on Gabor filter banks or wavelet transforms (Soltanian-Zadeh et al., 2004) rely on spatial frequency analysis of image regions, while as others use invariance properties of moments with respect to geometric transformations e.g., descriptors based on Zernike moments (Teague, 1980). A diagrammatic view of some these image descriptors is depicted in Figure 1.

**Figure 1:**
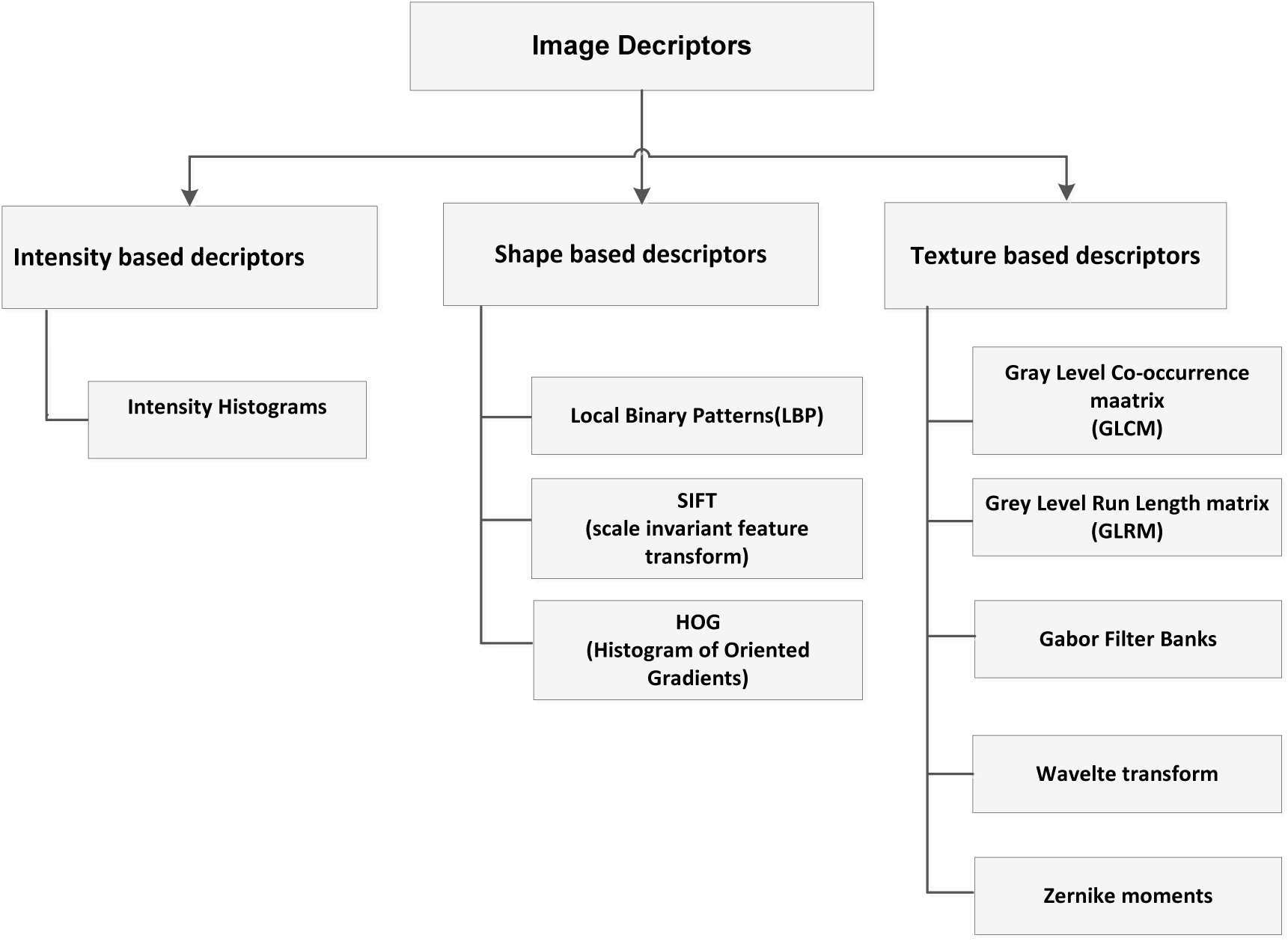
Different types of image descriptors.

Some of the important image descriptors based on the image property they are associated with are as follows: -

#### Based on intensity patterns

Intensity patterns are captured as a histogram of intensity values. The graph shows the frequency of each intensity value ranging from 0-255. For an 8-bit gray scale image having 256 different intensities, the histogram will display all values from 0-255 showing the distribution of pixels.

#### Based on shape patterns

- Shape based descriptors are characteristic of patterns that are related to image appearance or shape. Popular descriptors from this category include: -

- **SIFT (Scale invariant feature transform)**: SIFT transforms an image into a feature vector collection that is invariant to image rotation, translation, scale and robust to geometric distortions (Jurie & Schmid, 2004). SIFT algorithm works by detecting local extrema (maxima and minima) as key points in a scale space obtained by performing a difference of Gaussians on the input image. Once key points are detected, the algorithm performs a Taylor expansion of the scale space to obtain a more accurate location of key points. Thereafter, orientation is assigned to each key point to achieve invariance to image rotation (Lowe, 2004).
- **HOG (Histogram of gradients)**: HOG is an image descriptor used for detection and recognition of objects in image processing and computer vision (Tsai, 2010). The algorithm works on a localized image segment by counting the occurrences of gradient orientation.HOG descriptor algorithm works as follows: -

1. Partitions each image into cells in order to compute edge orientations for pixels within the cell.
2. Using gradient orientation, these cells are discretized into angular bins.
3. Each cell’s pixel contribution adds gradient (weighted) to its corresponding angular bin.
4. Cells that exist as neighbors are grouped as spatial regions called blocks. These blocks serve as a basis for grouping and normalization of histograms.
5. This normalized group represents the block histogram and a set of these block histograms represent the descriptor.HOG are very powerful descriptors and have been applied to object detection in computer vision (Dalal, 2005).
- **LBP (Local binary patterns)**: LBP is an image descriptor usually used for image classification in image processing or computer vision (Ojala et al., 2002). The algorithm works as follows:

1. An ROI is selected and divided into cells with 16x16 pixels each.
2. A comparison score of pixels being smaller or greater than the centre is represented as an 8 bit binary string which is later converted into decimal.
3. A count of frequency of each score is then outputted as 256 dimensional feature vector. The histogram is normalized and all histograms of the examined window are concatenated to obtain an LBP descriptor.
- *Based on texture patterns*: Texture is an important image property that helps in their characterization and recognition. Popular candidates from this category include GLCM (gray level co-occurrence matrix), GLRL (grey level run-length matrix), waveletes,curveletes and Gabor filter banks.
- **GLCM (Grey level co-occurrence matrix)**: GLCM characterises the texture of an image and populates a matrix of how often a pair of intensity values occur in an image. Once the matrix is defined, statistical measures such as homogeneity, entropy, contrast, average and correlation is computed (Haralick et al., 1973). These statistical measures have been shown to capture information of higher order from the image and are very robust in texture classification.
- **GLRL (Grey level run length matrix)**: A set of consecutive, collinear picture points having the same grey level value, constitutes a grey level run (Galloway, 1975). Number of picture points within the run determine the length of run. For a given image, we can compute a GLRL for runs having any given direction. The matrix element (*i*,*j*) specifies the frequency of *j* run lengths within the picture, in the given direction of points having grey level *i*.
- **GBF (Gabor filter banks)**: Gabor filter (Porat & Zeevi, 1988) is a linear bandpass filter used in image processing for feature extraction and edge detection. Their frequency and orientation representation corresponds to that of human visual system, and they have been found to be particularly appropriate for texture analysis, stereo disparity estimation and discrimination
- **DWT (Discrete wavelet transform)**: DWTs allow analysis of images into progressively finer octave bands. DWT’s are multi resolution analysis tools that enable to identify invisible patterns in the raw data. Using wavelets an image (2–D) approximation can be constructed that retains only desired features. There is a family of transforms based on wavelet packets that partition the frequency content of signals and images into progressively finer equal-width intervals. The application of wavelet and curvelet descriptors for mammographic analysis can be found in (Eltoukhy et al., 2010).

## 4 Computer Aided Diagnosis (CAD)

This section presents few application of computers and computer based analysis tools by citing some literature in the area. It gives a bird’s eye view of the role computers and machine intelligence can play in the area of computed based diagnostic descision making.

The analysis of medical images over the past several decades has become an important area for guided diagnosis and also as a tool to supplement the knowledge of healthcare professionals in general and radiologists in particular as they evaluate the physiological and anatomical state of a patient. Human analysis of biomedical images is usually subjective and qualitative, hence susceptible to errors. Therefore, computers along with machine intelligence algorithms can be used to encode the medical investigation via image analysis along with clinical information and devise procedures that can be applied consistently and objectively for such repetitive and routine diagnostic tasks (Doi, 2007).

CAD systems have their origin way back in 1963 when Lodwick, (1996) published his results on classification of pulmonary lesions on chest radiographs. But due to low computational power of computers and less advanced image processing techniques in those days, this area did not receive much attention until early 1990’s. As computer hardware became cheaper and sophisticated classification algorithms emerged in the domains of machine learning and artificial intelligence, the research focus on automated image analysis of biomedical images returned.

Recently, there has been a dramatic growth in the medical imaging studies and the discipline of radiology is witnessing a huge data explosion. The workload of radiologists has increased tremendously because of their limited number and thus the health care costs related to imaging are surging. In this scenario, computers become an implicit choice to process this huge data and help radiologists to make effective diagnostic decisions.

Modern day workstations are very powerful and state of the art computer programs for image processing do exist which can be exploited for diagnostic image analysis (Gonzalez, 2009). Computer vision techniques can be used to process biomedical images. A number of image descriptors discussed in the section(3) can be used to derive essential features from an ROI. These features are robust to geometrical transformation and invariant to scaling and can be used as feature vectors as input to highly sophisticated machine intelligence algorithms in order to distinguish normal patterns from abnormal ones.

With CAD radiologists use computer output as a second opinion while making a diagnostic decision. CAD has been a part of routine clinical work for detection of breast cancer on mammograms at many screening sites and hospitals across the world. Research efforts are underway to include more imagining modalities such as CT, MRI and PET scans as CAD schemes and implement them as a software / package in a PACS environment, a medical imaging framework for economical storage and convenient access to images from multiple modalities (Arenson, 1992).

## 5 Kernel Based Machine Learning

In this section we present basic concepts in machine learning such as, classification and the notion of cost functions. Thereafter, a detailed description about kernel methods, kernel trick and kernel matrices is given.

### 5.1 Basics in Machine Learning

#### Classification and Regression

Suppose there are *n* given objects ( *x_i_*) with 1 ≤ *i* ≤ *m* ∈ 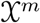 having labels ( *y_i_*) with 1 ≤ *i* ≤ *m* ∈ 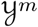. If their labels are finite, then we may be interested in classifying the given objects into different classes, assigning labels to unknown objects based on the examples observed a priori. However, in case the labels are continuous values, regression seems to be obvious choice. We can embed both of these tasks in a common framework where the job of classification is to predict discrete labels and that of regression continuous values (Alpaydin, 2014). The objects (*x_i_*)_1≤*i*≤*m*_ are termed as observations, inputs or simply as examples and the (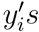) are termed as labels, targets or outputs. The set of all input pairs of (*x_i_*,*y_i_*)_1≤*i*≤*m*_ is known as the training set, whereas the test set is an unknown set of objects for which labels need to be predicted given the knowledge contained in the training set. Now given a function *f* defined on a set of objects ∈ 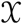, our main task is to generate correspondences (*x*_i_,*y*_i_) from the values contained in set 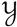, so that our model is able to generalize all possible correspondences of (*x_i_*,y*_i_*). The target for new objects in 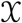 would then be *f*(*x*). By generalization, we mean for every *x* ∈ 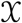 in the vicinity of an already observed *x_i_*, its target *f*(*x*) should also be very close to an already observed y_i_. This closeness in the space of 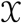 and 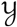 is vague if they are not vector spaces, therefore a precise similarity measure of inputs in 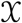 and penalties for predicting false targets in 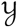 need to be defined accordingly. A graphical depiction of classification is shown in Figure 2.

**Figure 2:**
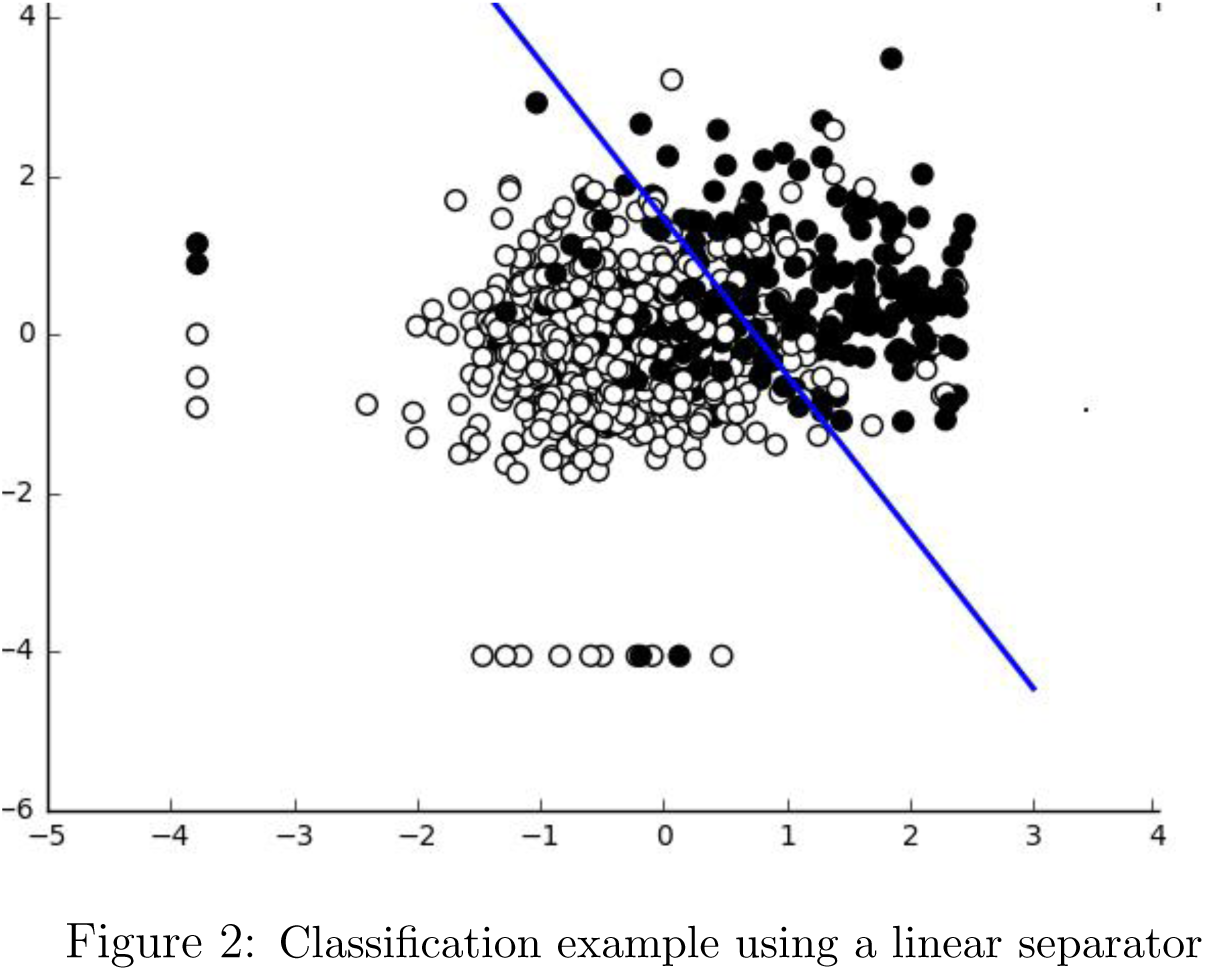
Classification example using a linear separator.

#### Cost Function

Measuring Euclidean distance in 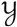 while dealing with regression is usually a convenient choice of similarity measure, but we can think of functions other than distances, provided they allow us to express penalties in case of wrong label assignment. Such functions, usually called the cost (C) or loss functions, account for all the penalties incurred (all costs) on all the mistakes made while searching for possible solutions *f* from the training data. We can have

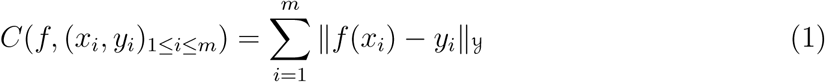

### 5.2 Similarity Measures and Features

Computing a similarity measure for objects (graphs, images, sequences, documents etc.) in 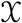 is not trivial, except that the space 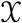 is chosen very carefully in which case defining a similarity measure is simply to compute the Euclidean distance between the points in vector space. Therefore, the challenge in learning is a careful choice N meaningful *features* Φ(*x_i_*) ( where Φ is a mapping from 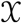 to 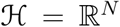. The notion is to design *features* Φ*_xi_* for each *x_i_* by selecting a more natural measure of similarity in the *feature space* (Zaki et al., 2014). Now selecting a feature map that is more natural or choosing a distance measure in 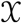 are equivalently hard problems. To find the best solution, i.e., a function *f* over a set of all possible functions *f*:, we can devise an optimization problem, such that it minimizes (2)

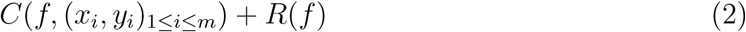
where *R*(*f*) is a regularization parameter that forces *f* to behave *smoothly* with respect to similarity measure in 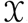.

### 5.3 Kernels

A kernel (Smola & Schölkopf, 1998) is defined as a measure of similarity on input 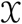 as

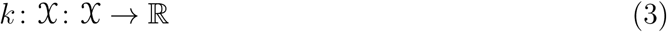

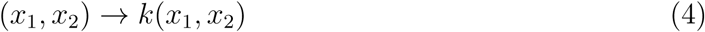
i.e., a function, whenever presented with two inputs *x*_1_ and *x*_2_ returns a real number *k*(*x_1_, x_2_*), a similarity score between them subject to the condition that it is symmetric for any two objects *x*_1_ and *x*_2_, i.e.,

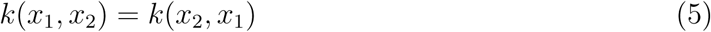

In choosing a similarity measure the standard Euclidean dot product in 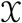 may not always be relevant or our input does not belong to vector space, in such circumstances a possibly meaningful feature set is available. We can then use this feature map as

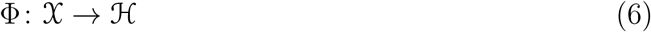

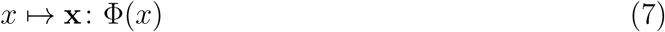

The bold face **x** is a vector representing our input in 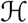. Φ is the feature that transforms the non-linear embedding in the input data and allows us to choose a similarity measure from a linear space.

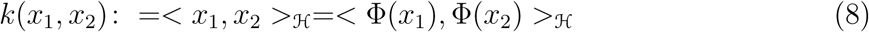

The flexibility in choosing Φ will enable us to design a number of machine learning algorithms and similarity measures. In the next subsection, we will see that in certain settings we don’t compute this explicit transformation (mapping via Φ) of our data, which is a convenient way of handling features in very high dimensional space.

### 5.4 Kernel Trick

The kernel trick is to show that the dot product Φ(*x_i_*)^*T*^ Φ(*x_i_*)in feature space can be replaced by a kernel function (Shawe-Taylor & Cristianini, 2004),

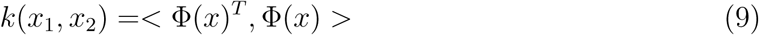
that can be efficiently computed in the input space. Geometrically we can interpret the input data through the definition of *k*,

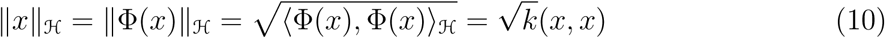
gives us the length of 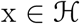. *k*(*x*_1_, *x*_2_) can similarly compute the cosine of angle between *x*_1_ and *x*_2_ provided they are normalized to a unit vector. Distance between two vectors can be computed as the length of their difference,

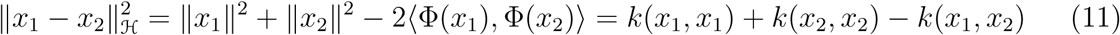

Now considering any such *k*(*x*_1_, *x*_2_) and without having to perform the mapping Φ explicitly, we can compute all the attributes viz. lengths, distance and angles using *k* only. The power of kernel methods framework lies in detecting any non-linear patterns in input data using linear algebra and analytical geometry, this is the kernel trick.

### 5.5 Kernel Matrix

Biomedical images are a rich source of structural and functional information of the living systems under study. This information can be extracted in the form of image descriptors as described in section 3. Each dataset is transformed into a symmetric positive semi-definite kernel matrix by means of a kernel function,that is a real valued *k* (*x*_1_, *x*_2_) satisfying *k* (*x*_1_, *x*_2_) = *k* (*x*_2_, *x*_1_) for any two objects *x*_l_ and *x*_2_ and positive semi-definite i.e., to say 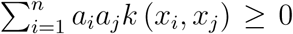 for any integer n, set of objects (*n* = *x*_l_…˙. *x*_n_) and any set of real numbers (*a*_l_…˙.*a*_*n*_) (Charpiat, 2015).

Kernel functions provide a coherent representation and a mathematical framework for the input data and represent the object features via their pairwise similarity values comprising the n × n *kernel matrix*,defined as.

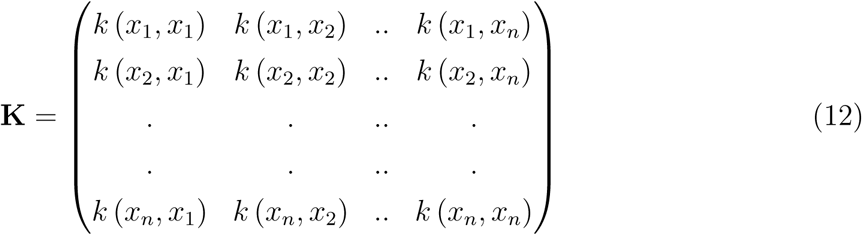

Kernel methods present a modular approach to pattern analysis (Shawe-Taylor & Cristianini, 2004). An algorithmic procedure is devised together with a kernel function that performs an inner product on the inputs in a feature space. This algorithm is more generic and can work for any kernel and hence for any data domain. The kernel part is data specific that offers an elegant and flexible approach to design learning systems, that can easily operate in very high dimensional space. It is a modular framework, where modules are combined together to obtain complex learning systems. Some examples of commonly used kernels are:

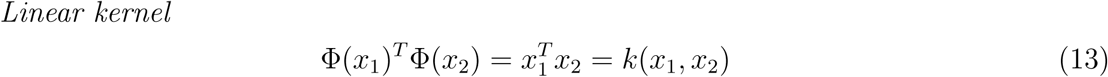

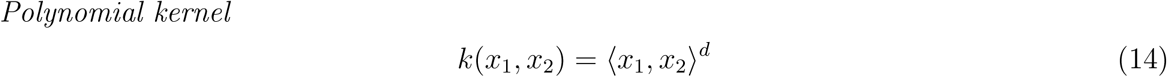

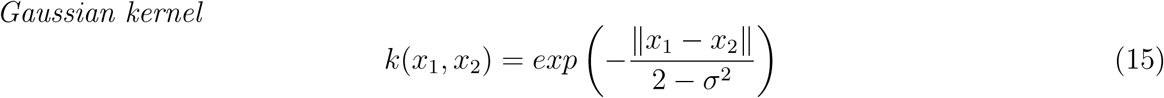

For suitable values of *d* and *σ*, the similarity measure *k*(*x*_1_, *x*_2_) between *x*_1_ and *x*_2_ is always positive with is maximum at *x*_1_ = *x*_2_. All points *x* have same unit norm (since *k*(*x*_1_, *x*_2_) = 1∀ _*x*_) suggesting that images of all points in x lie on the unit sphere in 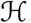.

## 6 Multiple Kernel Learning Model

This section cover the multiple kernel learning (MKL) paradigm in machine learning and optimization of cost function using SDP (semidefinite programming).

Exploiting this modular approach to learning system design, multiple kernel learning (MKL) is a paradigm shift from traditional single feature based learning and offers an advantage of combining multiple features of objects such as images, documents, videos etc.,as different kernels (Sonnenburg et al., 2006). This information can be fed as an ensemble into an MKL learning algorithm as a combined kernel matrix for classification or regression tasks on unknown data. The basic algebraic operations of addition, multiplication and exponentiation when performed in combining multiple kernel matrices preserves the positive semi-definite property and enable the use of powerful kernel algebra. A new kernel can be defined using *k_1_* and *k_2_* with their corresponding embeddings Φ_1_(*x*)and Φ_2_(*x*). This resultant kernel is

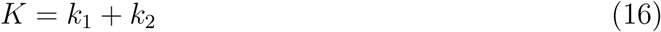
with the new induced embedding

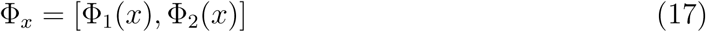

Given a kernel set *K* = {*k*_1_, *k*_2_,…., *k_m_*}, an affine combination of m parametrized kernels can be formed as given below: -

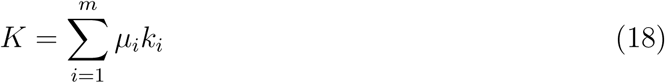
subject to the constraint that *μ_i_* (weights) are positive i,e. *μ_i_* ≥ 0, *i* = 1……˙.*m* A kernel based statistical classifier such as SVM induces a margin in feature space, separating the two classes using a linear discriminant. In order to find this linear discriminant, an optimization problem needs to be solved, known as a quadratic program (QP). A Quadratic programs is a form of convex optimization problem, which are easily solvable. On the basis of this margin, SVM algorithms are classified into hard margin, 1-norm soft margin and 2-norm soft margin SVM. We will introduce 1-norm soft margin SVM here, for further literature on SVM algorithms see Smola & Schölkopf, 1998.

**1-Norm Soft Margin SVM**

An SVM algorithm produces a linear separator in the feature space.

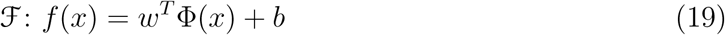
where 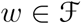 and 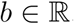. Given an input data set 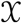 with label set 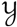,set of all correspondences from these two sets S_n_ = {(*x*_1_,*y*_1_),…˙., (*x*_n_,*y*_n_)} is a labeled set. A 1-norm soft margin support vector machine selects an optimal discriminant, optimizing with respect to *w*&*b* maximizing the margin between two classes and allowing misclassification to some degree, hence the name soft margin.

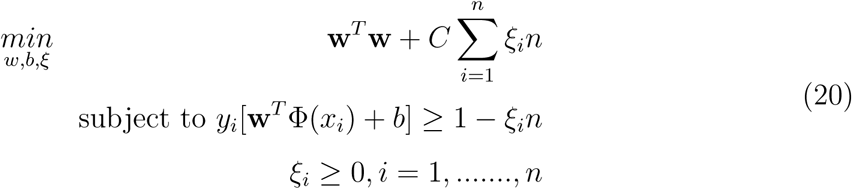

Parameter *C* is used for regularization maintaining a trade off between error and the decision boundary (margin), ξ is a *slack variable*, 0 < ξ < 1 which means that data point lies somewhere between the margin and correct side of the hyperplane and if ξ > 1, it means that data point is misclassified. By taking the dual of equation(2), the weight vector can be expressed as 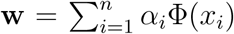 The values of *α_i_*, called the support values, these are considered to be solutions for the following dual problem,

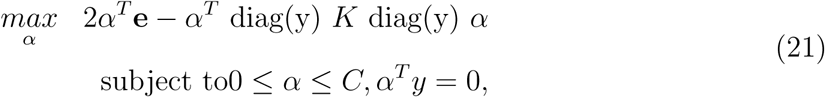

Where diag(**y**) is a diagonal matrix whose entries are elements in *y* = (*y*_l_,*y*_2_…….*y*_n_). An unknown data item (*x_new_*) can be assigned a label by computing the linear function,

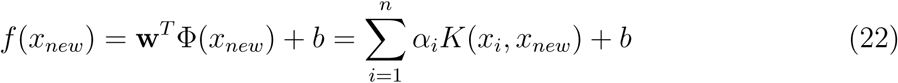

Now depending upon the value returned by *f*(*x*_*new*_), we can classify *x_new_* either bearing a class lable of +1 or −1. SVM has been successfully applied to image analysis and classification based on intensity histograms in (Chapelle, 1999)

### 6.1 MKL optimization using SDP

Multiple kernel learning (MKL) is based on convex combinations of arbitrary kernels over potentially different domains and can be solved using semidefinite programming (SDP) optimization techniques. Lanckriet et al., (2004a) have shown that the classification performance for a fixed trace of kernel matrix is bounded by a function, whose optimum is achieved in Equation(21), a smaller value guarantees better performance and vice versa. In SVM, we use a single kernel matrix *K*. We can formulate an extended version of Equation(21) by parameterizing *K* in order to learn an optimizied kernel matrix. Taking additional constraints such as,trace and positive semi-definiteness in to consideration for Equation(19), and minimizing with respect to parameters *μ_i_* Equation(21) can be rewritten as:

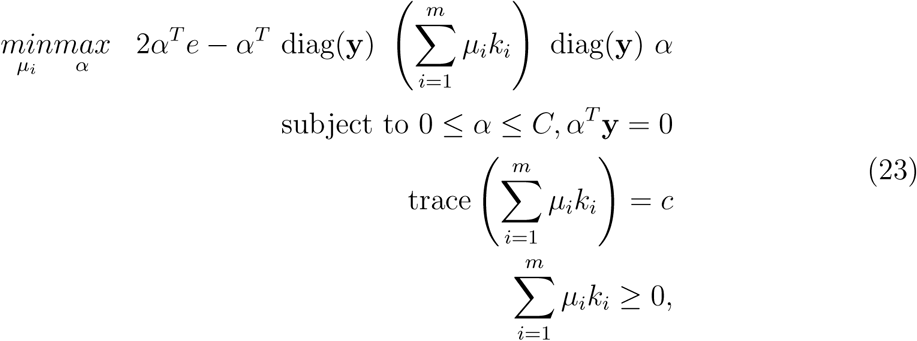

Where c is a constant. Now, if we take a Lagrangian dual of the problem, it has been show in (Lanckriet et al., 2004a) that finding optimum *μ_i_* and *α_i_* reduce this Lagrangian to a semi-definite program (SDP), a form of convex optimization problem.

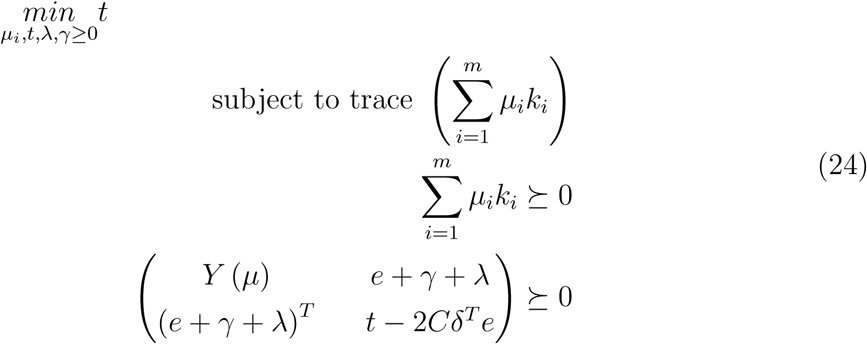

Where

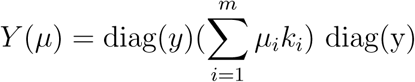

Semidefinite programming presents a general case of LP (linear programming), where the linear matrix inequalities (LMI) replace the scalar inequalities of LP’s. Solution for both LP and SDP can be obtained via interior point algorithms (Nesterov & Nemirovskii, 1994) as both of them are instances of convex optimization problems. For the Multiple kernel learning problem we consider weights *μ_i_* ≥ 0 and *K*_*i*_’s are PSD matrices. Equation(24) SDP can be reduced to quadratically constrained quadratic program (QCQP) that can be solved efficiently as shown in Lanckriet et al., (2004a):

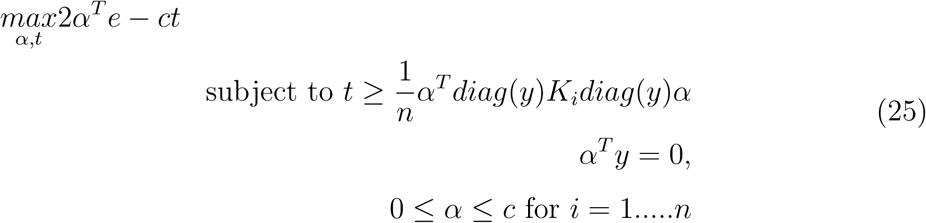

The solution to our problem of classification lies in solving the above QCQP. By solving this quadratic program we are in a position to find a combination of kernel matrices that are adaptive, robust and flexible in handling heterogeneous sources of data. Our classification task is achieved based on information encoded in multiple kernels and the output weights reflect the relative contribution of each information source. MKL has been successfully applied in the domain of bioinformatics for functional classification of genes (Lanckriet et al., 2004b), protein classifications and inferring protein-protein interactions in yeast and is a suitable candidate for analyzing biomedical images. (Chapelle, 1999)

## 7 Multiple Kernel Learning for Biomedical Image Analysis

In this section we present a framework based on **openCV** (Open source computer vision) technology and **ShOGUN** machine learning toolbox for classifying mammograms using HOG,SIFT and LBP descriptors. We will briefly describe these tools here before presenting experimental steps performed on mammograms of 150 patients from **MIAS** (Mammographic Image Analysis Society) database. A graphical representation of the framework that has been implemented in python using shogun and openCV is depicted below in Figure 3.

**Figure 3:**
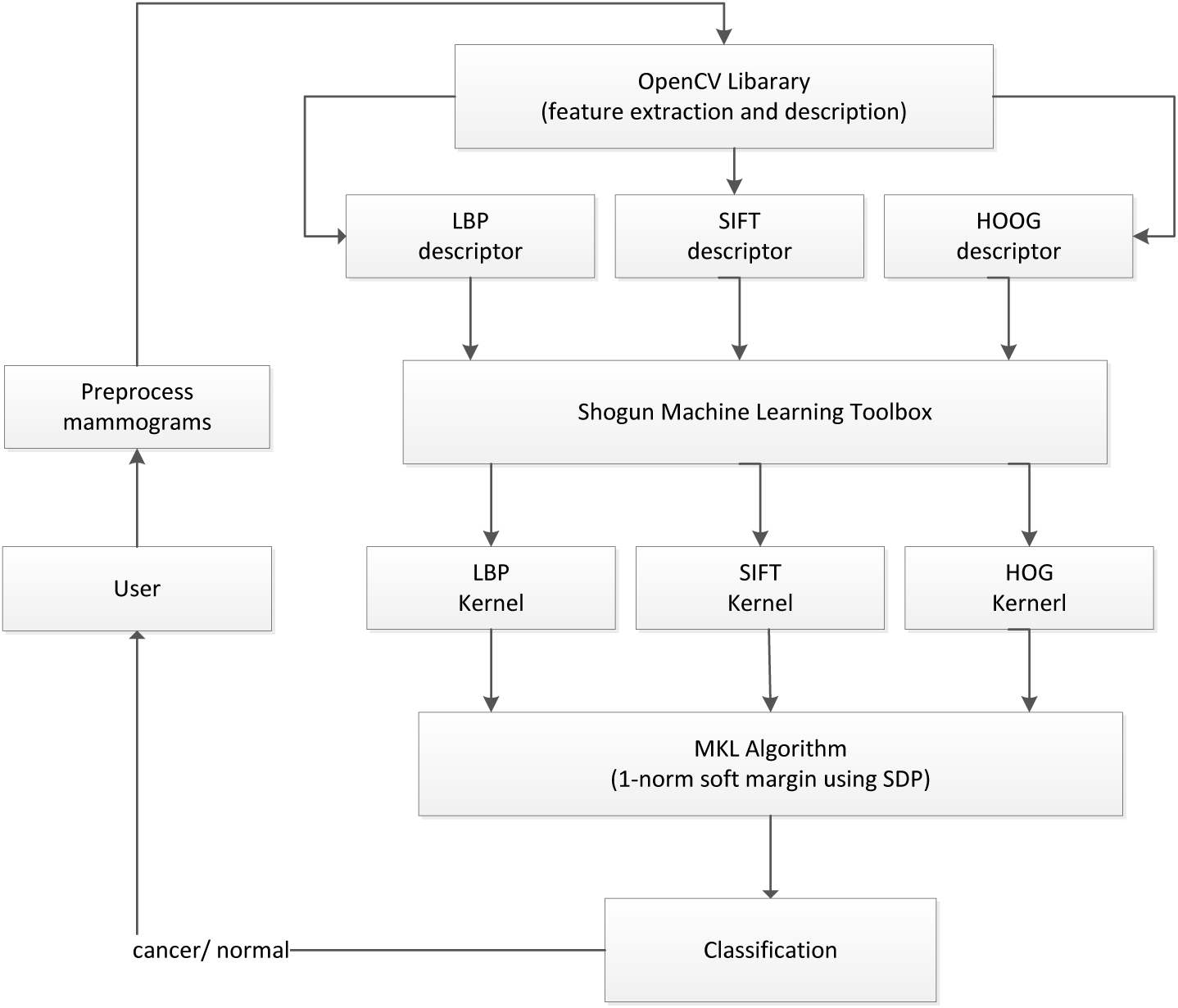
Diagrammatic representation of biomedical image analysis using MKL.

Biomedical images alone may not serve as direct indicators of the pathological symptoms of a patient. In fact medical image analysis done in isolation may lack a direct relationship to any patient illness. However, the information captured by various imaging modalities can provide important clues to radiologists and other health care professionals while evaluating the state of a patient. As described in section 3, different types of image descriptors capture different information, e.g, shape, texture moments and intensity, we can extract these features from medical images, process them and use computer algorithms to predict the relationship between image features and patient pathology. The output from this system along with expert knowledge will equip health care evaluators with better diagnostic decisions and therefore a better treatment for the patient.

### 7.1 OPENCV (Open Source Computer Vision)

OpenCV is an open source machine learning and computer vision library. The OpenCV platform provides a unified and efficient infrastructure for applications in the domain of pattern analysis and computer vision. OpenCV is BSD licensed with more than 2500 optimized software algorithms for machine learning, image processing and computer vision, which includes a comprehensive set of both classic and state-of-the-art algorithms from these domains. These algorithms can be freely used for academic as well as commercial purposes and are used to detect variety of objects, textures from images and are even able to classify movements in videos, follow eye movements recognize scenery, etc. from videos. OpenCV libraries are available for all platforms viz, Windows, Linux, Mac OS and Android. With basic implementation in C++, OpenCV provides interfaces to other high level languages, such as Java, Python and Matlab etc. We have used OpenCV libraries here to extract descriptors from mammographic images, for detailed documentation refer(OPENCV, n.d.)

### 7.2 Shogun Machine Learning Toolbox

The Shogun machine learning toolbox offers plenty of algorithms and methods to solve complex problems, such a classification, regression, dimensionality reduction and clustering etc. The toolbox provides increased flexibility to work with data representations, algorithmic classes and other problem solving tools like CPLEX, LPSOLVE,MOSEK etc.

Shogun is implemented in C++ and provides interfaces to many modern day high level languages and tools, such as Java, Python, Octave, Matlab and R etc. Blending modern software architecture with state-of-the-art algorithmic implementation, shoigun platform provides a rich library of tools and methods for solving large scale machine learning problems, even with modest compute infrastructure.

Shogun is licensed under GPL and has been a tool of choice in the areas of large-scale kernel based learning and bioinformatic. In this tutorial we use Shogun MKL classes for classification of mammographic data. For further details see (Sonnenburg et al., 2010)

### 7.3 Case Study

This subsection describes the use of above framework for classifying mammograms. Following a brief description of the database, we describe experimental steps performed and the results obtained thereafter.

#### 7.3.1 Dataset

A collection of 300 mammograms (1024×1024 in resolution) in portable gray map format (.pgm) from MIAS (Mammographic image Analysis society) mini database is downloaded from PEIPA project website (Pilot European Image Processing Archive). The mammograms are then subjected to feature extraction process using OpenCV tools for SIFT,HOG, and LBP descriptors etc. Once the features are obtained, each feature set is divided into training and testing set in the ratio of 70:30. Feature such as, SIFT and LBP are further processed to obtain feature vectors. Feature vectors for each set are stacked vertically to form datasets in shogun format.

#### 7.3.2 Experiment

Because we have chosen kernel based learning, it is important to transform each dataset of features into a kernel matrix. We choose Gaussian kernel with an initial width =10 for our MKL approach, Gaussian kernels perform well with the vectorial input (Smola & Schölkopf, 1998).

Kernels for all the features, viz. SIFT,HOG and LBP are joined together using *CombinedFeature()* class of Shogun toolbox. This fused kernel along with data labels are then passed to an instance of *MKL()* class. Initially, a training set is passed as input. The algorithm is evaluated after passing the test data. The results obtained are compared with other binary classifiers, viz. KNN,SVM and Naive Bayes, which are run individually on all features.Table 2 shows the results of all the runs from all the classifiers.

**Table 2:**
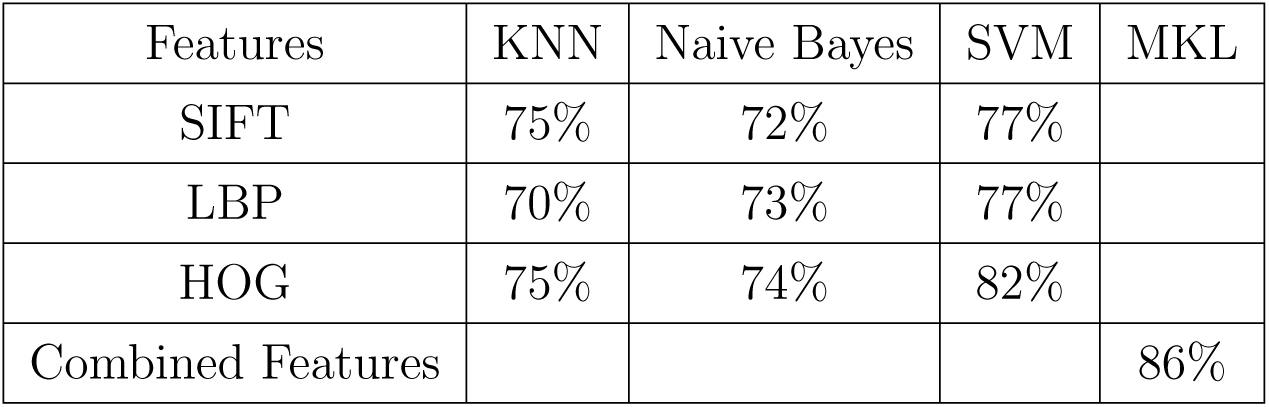
Show classification results of various classifiers using SIFT,HOG,LBP features. MKL shows a marked improvement as it takes contribution of all.

| Features | KNN | Naive Bayes | SVM | MKL |
| --- | --- | --- | --- | --- |
| SIFT | 75% | 72% | 77% |  |
| LBP | 70% | 73% | 77% |  |
| HOG | 75% | 74% | 82% |  |
| Combined Features |  |  |  | 86% |

To boost the learning process of MKL algorithm, k=5 fold cross validation is performed using the input data, this yields an optimum kernel width of 3.17 with a performance boost of 2% in our *1-norm* soft margin MKL binary classifier.

## 8 Discussion & Conclusion

After covering feature extraction, image description and a tutorial on kenel methods and multiple kernel learning model(MKL), we have a presented a framework for integrating multiple features to enhance the prediction accuracy of a binary classifier using MKL. The performance of the system can be furthe improved upon if the kernel machine are trained with large amounts of input data. From Table 2, it is evident that combined features reveal more information to the classifier as compared to feeding them individually. This experiment is a small effort in highlighting the power of MKL paradigm for improved classification using multiple feature sets against other classifiers that work on a single feature set. We hope this tutorial wakes up to reader to apply the MKL framework to a larger image dataset and to different types of biomedical images where the raw data and clinical information is available.

Modern day machine learning approaches to biomedical image analysis rely heavily on the quality of features extracted from the medical images and their relevance to the pathological symptoms under investigation. Since body subsystems possess features comprising of shape, texture,density and signals, it is better to capture as much information as possible from the diagnostic imaging in order to better understand the state of a patient. A mathematical framework to integrate these different information sources, such as kernel methods and multiple kernel learning will contribute immensely in the design of CAD systems, which are robust to image distortions. Integrating such a system in a PACS environment can revolutionize the diagnostic image analysis and improve the productivity of healthcare professionals both interms of better diagnosis and better treatment.

Although lot of research is going on in the field of biomedical image analysis, but the challenges to develop efficient algorithms for advanced imaging modalities and to improve the accuracy of existing ones are opening new vistas in this domain. A number of GRAND challenges are organized in order to attract the attention of research community, both from academia and industry or solving complex image analysis problems. Some of these challenges include, improving accuracy in digital mammography analysis for early detection of breast cancer, automatic segmentation of PET images for delineation of tumors, diagnostic classification of clinically significant prostrate lesions, developing algorithms which can accurately identify types of cervical cancers in women and detecting tumor proliferation across various tissues etc. Besides these, there are a number of other open research challenges in the field that need an immediate attention of the researchers from diverse domains, such as biophysics, computer science, radiology, computer vision, machine learning and machine intelligence.

## ABBREVIATIONS

BSD: Berkely software distribution
CAD: Computer aided diagnosis
CT: Computed Tomography
DWT: Discrete wavelet transform
GFB: Gabor lter bank
GLCM: Grey level co-occurenece matrix
GLRLM: Grey level run length matrix
HOG: Histogram of oriented gradients
KNN: K nearest neighbor
LBP: Local binary patterns
LP: Linear programming
MIAS: Mammographic Imaging Analysis Society
MKL: Multiple kernel learning
MRI: Magnetic Resonance Imaging
PACS: Picture archive communication system
PEIPA: Pilot European Image Processing Archive
PET: Positron Emission Tomography
ROI: Region of interest
SDP: Semi-denite programming
SEM: Scanning electronic microscope
SIFT: Scale invariant transform
SPECT: Single photon Emission Computed Tomography
SVM: Support vector machine
TEM: Transmission electronic microscope

